# Circumpolar diversification of the *Ixodes uriae* tick virome

**DOI:** 10.1101/2020.05.04.075515

**Authors:** John H.-O. Pettersson, Patrik Ellström, Jiaxin Ling, Ingela Nilsson, Sven Bergström, Daniel González-Acuña, Björn Olsen, Edward C. Holmes

## Abstract

Ticks (order: Ixodida) are a highly diverse and ecologically important group of ectoparasitic blood-feeding organisms. One such species, the seabird tick (*Ixodes uriae*), is widely distributed around the circumpolar regions of the northern and southern hemispheres. It has been suggested that *Ix. uriae* spread from the southern to the northern circumpolar region millions of years ago and has remained isolated in these regions ever since. Such a profound biographic subdivision provides a unique opportunity to determine whether viruses associated with ticks exhibit the same evolutionary patterns as their hosts. To test this, we collected *Ix. uriae* specimens near a Gentoo penguin (*Pygoscelis papua*) colony at Neko harbour, Antarctica, and from migratory birds - the Razorbill (*Alca torda*) and the Common murre (*Uria aalge*) - on Bonden island, northern Sweden. Through meta-transcriptomic next- generation sequencing we identified 16 RNA viruses, seven of which were novel. Notably, we detected the same species, Ronne virus, and two closely related species, Bonden virus and Piguzov virus, in both hemispheres indicating that there have been at least two cross- circumpolar dispersal events. Similarly, we identified viruses discovered previously in other locations several decades ago, including Gadgets Gully virus, Taggert virus and Okhotskiy virus. By identifying the same or closely related viruses in geographically disjunct sampling locations we therefore provide evidence for virus dispersal within and between the circumpolar regions. In marked contrast, our phylogenetic analysis revealed no movement of the *Ix. uriae* hosts between the same locations. Combined, these data suggest that migratory birds are responsible for the movement of viruses at both the local and global scales.

**Author summary/Importance:** As host populations diverge, so may those microorganisms, including viruses, that are dependent on those hosts. To examine this key issue in host-microbial evolution we compared the co-phylogenies of the seabird tick, *Ixodes uriae*, and their RNA viruses sampled from the far northern and southern hemispheres. Despite the huge geographic distance between them, phylogeographic analysis reveals that the same viruses were found both within and between the northern and southern circumpolar regions, most likely reflecting transfer by virus-infected migratory birds. In contrast, genomic data suggested that the *Ix. uriae* populations were phylogenetically distinct between the northern and southern hemispheres. This work emphasises the importance of migratory birds and ticks as vectors and sources of virus dispersal and introduction at both the local and global scales.

## Introduction

Following the physical separation of a population into geographically isolated sub-populations (i.e. vicariance) genetic changes unique to each sub-population will accumulate. Given a sufficient period of time, such process may result in marked genetic separation. By combining phylogenetic and geographical information - that is, phylogeography [1,2] - it is possible to infer the spatial and evolutionary relationships among such subdivided populations. The analyses of these populations may include inferences on the direction of dispersal between subpopulations and if there have been multiple introductions into a particular geographic region. A recently colonised area is expected to exhibit less genetic diversity than the source population [1,2].

As host populations diverge, so will any microorganisms, including viruses, that are dependent on their host(s). Accordingly, analysis of genome sequence data from these microorganisms can provide additional, and sometimes more detailed, information about the evolutionary history and demography of the host species [3,4]. Resolution of the patterns and processes of host-pathogen co-divergence is particularly strong in the case of RNA viruses in which mutational changes accumulate much faster than in their hosts [4]. For example, analysis of the phylogeny of feline immunodeficiency virus (FIV) provided important information on the recent population history and demography of its feline host, the cougar *Puma concolor*, that was not apparent in host genetic data [5].

Ticks (order Ixodida) are among the most diverse groups of ectoparasites. There are close to 900 species of both soft- and hard-bodied ticks within the Ixodida [6,7], of which the genus *Ixodes* is the most species rich group with nearly 250 species [6,8–10]. Within this genus, *Ix. uriae* is the only known tick with a circumpolar distribution in both the northern and southern hemispheres. This species parasitizes close to 100 different vertebrate species, the majority of which are seabirds that breed in dense colonies [11]. In the northern hemisphere, the most commonly recorded hosts are birds of the order Charadriiformes, mainly *Alcidae* and *Laridae*, and in the southern hemisphere they are mainly species of the Spheniciformes and Procellariiformes [11–14]. Like most hard ticks, *Ix. uriae* has three active life-stages (larva, nymph and adult) whose questing behaviour is most prevalent during the summer months with a peak during June–July in the northern hemisphere and December–January in the southern hemisphere [15–18]. Each active stage takes a single blood-meal from a host during 3–12 days depending on the tick’s life-stage. The duration of the life cycle depends on environmental temperatures and may last from 3 to 7 years, among the longest seen in ticks [13,15,19,20]. Importantly, and perhaps as a consequence of its host and habitat adaptation, *Ix. uriae* has a temperature tolerance to as low as −30°C and as high as +40°C [21], forming aggregations in moist rocky microhabitats [17,21]. *Ix. uriae* is a well-known vector of multiple different viruses and bacteria, including *Borrelia burgdorferi* sensu lato [22,23], the agent of Lyme disease, and Gadgets Gully virus [24,25] amongst others (reviewed in [11]). However, although some viruses and bacteria may have zoonotic potential and humans are bitten by *Ix. uriae*, there is limited evidence for human disease associated with *Ix. uriae* transmission [11].

Data from both mitochondrial and nuclear genes suggest that *Ix. uriae* diverged from its most recent common ancestor, *Ix. holocyclus*, approximately 91 million years ago, and that the *Ix. uriae* species complex shared a common ancestor some 22 million years ago [26]. Subsequently, *Ix. uriae* was introduced, possibly twice, into the northern hemisphere from the likely ancestral Australasian population approximately 10 million years ago [26]. Thereafter, both the southern and northern populations diversified into geographically structured subpopulations with no evidence of dispersal between them [26]. However, as some birds can migrate long distances, it is important to determine whether there has been any recent viral dispersal between the two polar regions and if there is any gene flow between the two *Ix. uriae* sub-populations. Here, by comparing the viromes of *Ix. uriae* collected from seabirds – the Common murres’ (*Uria aalge*) and Razorbills’ (*Alca torda*) – from the northern hemisphere and from around a penguin (*Pygoscelis papua*) colony in the southern hemisphere, we investigated whether there has been virus dispersal either within or between the northern and southern hemispheres.

## Results

In total, we generated 16 RNA sequencing libraries from 33 ticks, all of which were engorged adult female *Ix. uriae* individuals: 10 libraries using two tick individuals from the southern hemisphere, and six libraries from the northern hemisphere comprising five with two tick individuals and one with three tick individuals, (Supplementary Table S1). These libraries were sequenced to a high depth and assembled *de novo*. Across the libraries as a whole we identified 16 RNA viruses, seven of which were novel based on RNA-dependent RNA polymerase (RdRp) sequence similarity. The viruses identified belong to the following orders/families: *Bunyavirales* (N = 5), *Hepeviridae* (N = 2), *Flavivirus* (genus) (N = 1), *Mononegavirales* (N = 1), *Orthomyxoviridae* (N = 2), *Picornaviridae* (N = 1), *Reoviridae* (N = 2) and *Tombusviridae* (N = 2) (Figure 1).

**Figure 1.**
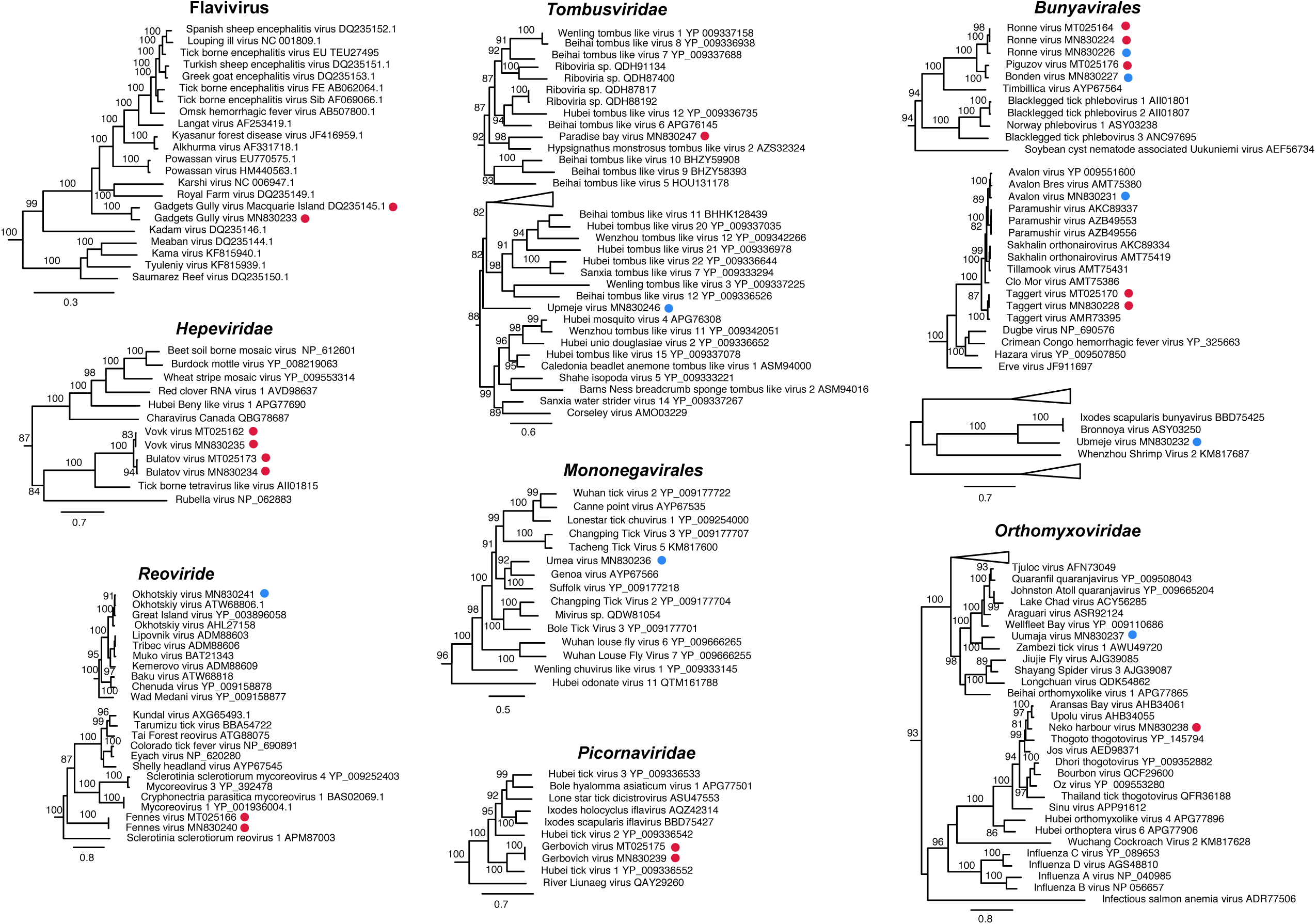
Phylogenetic analysis of all the viruses identified here within their respective virus groups, including representative publicly available viruses. Viruses identified in the current and a related study (27) are indicated, with red and blue circles for viruses identified from the southern and northern circumpolar regions, respectively. Numbers on branches indicate SH support (only branches with SH support ≥80% are indicated) and branch lengths are scaled according to the number of amino acid substitutions per site. All phylogenetic trees were mid-point rooted for clarity only.

The individual viruses discovered were found at abundance levels ranging from 3 to 10,203 reads per million and the total viral abundance per library, approximated from all virus RdRp reads mapped in positive libraries, varied between 14–12,257 reads per million. Correspondingly, the abundance of *Ix. uriae*, approximated via the COX1 gene, varied between 1,359–8,101 reads per million (Table 1). Four viruses - Gadgets Gully virus (*Flavivirus*), Taggert virus (*Bunyavirales*), Neko harbour virus (*Orthomyxoviridae*) and Upmeje virus (*Tombusviridae*) - were found to be highly abundant (i.e. abundance levels above 1,000 reads per million or more than 0.1% of the total number of reads per library). Indeed, Gadgets Gully virus, Taggert virus and Upmeje virus were more abundant than the host COX1 abundance (Table 1), reaching 1,704, 1,768 and 10,203 reads per million, respectively, and suggesting that they are tick-associated viruses.

**Table 1.** Presence and estimation of virus abundance across libraries

| Virus name | *Ixodes uriae*, Antarctica |  |  |  |  |  |  |  |  |  | *Ixodes uriae*, northern Sweden |  |  |  |  |  | IH 0.1% |
| --- | --- | --- | --- | --- | --- | --- | --- | --- | --- | --- | --- | --- | --- | --- | --- | --- | --- |
|  | IU1 | IU2 | IU3 | IU4 | IU5 | IU6 | IU7 | IU8 | IU9 | IU10 | IU15 | IU16 | IU17 | IU18 | IU19 | IU20 |  |
| Ronne virus | 14 | 24 | 19 | 16 | 21 | 22 | 4 | 0 | 4 | 7 | 1 | 3 | 6 | 1 | 1 | 1 | 1 |
| Ronne virus | 1 | 3 | 2 | 2 | 2 | 2 | 1 | 0 | 0 | 1 | 10 | 24 | 69 | 7 | 9 | 3 | 3 |
| Bonden virus | 0 | 1 | 0 | 0 | 1 | 0 | 1 | 0 | 0 | 0 | 7 | 13 | 45 | 3 | 4 | 1 | 2 |
| Taggert virus | 1 | 4 | 1 | 5 | 1 | 971 | 2175 | 0 | 1768 | 1 | 5 | 1 | 1 | 1 | 1 | 1 | 103 |
| Avalon virus | 0 | 0 | 0 | 1 | 0 | 11 | 25 | 0 | 21 | 0 | 312 | 0 | 0 | 0 | 0 | 0 | 15 |
| Ubmeje virus | 0 | 0 | 0 | 0 | 0 | 0 | 0 | 0 | 0 | 0 | 17 | 203 | 1 | 0 | 0 | 41 | 13 |
| Gadgets Gully virus | 0 | 0 | 0 | 3 | 0 | 0 | 0 | 0 | 1704 | 0 | 0 | 0 | 1 | 0 | 0 | 0 | 75 |
| Bulatov virus | 0 | 70 | 88 | 101 | 39 | 0 | 35 | 8 | 12 | 92 | 0 | 0 | 0 | 0 | 0 | 0 | 5 |
| Vovk virus | 0 | 14 | 13 | 136 | 24 | 0 | 236 | 3 | 90 | 11 | 0 | 0 | 0 | 0 | 0 | 0 | 4 |
| Umea virus | 1 | 0 | 1 | 0 | 0 | 1 | 0 | 0 | 0 | 1 | 73 | 506 | 1438 | 135 | 92 | 15 | 66 |
| Uumaja virus | 1 | 0 | 0 | 1 | 0 | 0 | 0 | 0 | 0 | 1 | 59 | 233 | 502 | 44 | 54 | 80 | 23 |
| Neke harbour virus | 0 | 0 | 1 | 0 | 2008 | 0 | 0 | 0 | 2 | 85 | 0 | 1 | 0 | 0 | 0 | 1 | 95 |
| Gerbovich virus | 0 | 369 | 3 | 0 | 552 | 0 | 0 | 0 | 3 | 0 | 0 | 0 | 0 | 0 | 0 | 0 | 26 |
| Fennes virus | 0 | 0 | 12 | 61 | 0 | 14 | 0 | 0 | 172 | 0 | 0 | 0 | 0 | 0 | 0 | 0 | 8 |
| Okhotskiy virus | 0 | 0 | 0 | 0 | 0 | 0 | 0 | 0 | 0 | 0 | 0 | 296 | 0 | 0 | 0 | 0 | 19 |
| Upmeje virus | 3 | 2 | 2 | 2 | 2 | 1 | 1 | 0 | 2 | 1 | 2 | 2 | 10203 | 2 | 573 | 661 | 469 |
| Paradise bay virus | 4 | 0 | 0 | 0 | 0 | 0 | 0 | 115 | 0 | 0 | 0 | 0 | 0 | 0 | 0 | 0 | 48 |
| *Ix. uriae* COX1 | 1548 | 2555 | 2967 | 1359 | 3520 | 2824 | 3747 | 2598 | 1505 | 1807 | 8101 | 4678 | 1964 | 3179 | 4819 | 4580 |  |
| Total virus*** | 14 | 476 | 132 | 315 | 2645 | 1006 | 2449 | 1231 | 3750 | 110 | 478 | 1275 | 12257 | 189 | 732 | 765 |  |

IH 0.1% refers to the index-hopping cut-off in relation to the most abundant library.
\* Values in bold indicate libraries with an abundance above 1000 reads per million.
\*\* Libraries in bold within a square indicates abundance above that of the host.
\*\*\* Total virus abundance for a single library as reads per million.

### Circumpolar virome comparison

Of the 16 viruses identified, nine were found in *Ix. uriae* ticks sampled from a Gentoo penguin (*Pygoscelis papua*) colony at Neko harbour, Antarctica, and seven were found from *Ix. uriae* ticks collected from Razorbill (*Alca torda*) and Common murre (*Uria aalge*) seabirds on Bonden island in the Gulf of Bothnia, northern Sweden (Figure 1, Table 1, Supplementary table 1). The geographical separation of the different virus species’ was confirmed when there was no overlap of individual virus contigs between the two sampling sites (i.e. a particular virus or virus variant was only found in the northern or southern hemisphere sampling site, but not both) (Table 1). Indeed, the majority of all viruses were found either in the northern or southern hemisphere *Ix. uriae* sequence libraries, but not both (Table 1, Supplementary table S2). Notably, however, we identified two variants of Ronne virus (*Bunyavirales*) in both Antarctica and northern Sweden (Figure 1, Table 1). Ronne virus has been previously identified from ticks in Antarctica [27]. One variant of Ronne virus identified here, also sampled from Antarctica, was highly similar (99.9% amino acid similarity; 99.7% nucleotide similarity) to that previously identified, whereas the second variant, recovered from ticks collected from the north of Sweden, was more divergent in sequence (93.2% amino acid similarity; 80.3% nucleotide similarity). Such genomic similarity is indicative of a relatively recent dispersal event between the northern and southern tick populations, although the direction of migration remains to be determined. Similarly, we identified a novel bunyavirus, Bonden virus, from northern Sweden that is the closest relative (90.3% amino acid similarity; 78.8% nucleotide similarity) of Piguzov virus identified from *Ix. uriae* in Antarctica (Figure 1) [27]. Although it is unclear when these two viruses diverged, they do point to an historical dispersal event between the two poles.

It was notable was that we identified both Taggert virus (*Bunyaviridae*) and Gadgets Gully virus (*Flaviviridae*) from the Neko harbour sampling site. Both these viruses have previously been found on Macquarie island, south-east of the Australian continent [24,25,28]. From northern Sweden we identified Avalon virus (*Bunyaviridae*) and Okhotskiy virus (*Reoviridae*), previously isolated from *Ix. uriae* from the Great island, Newfoundland, Canada, and from islands around the sea of Okhotsk, respectively [29–31]. The finding of Avalon virus, Gadgets Gully virus, Okhotskiy virus and Taggert virus at these new locations again suggests that there has been widespread geographical dispersal of viruses within each hemisphere.

Less clear is whether the divergence of these variants justifies their classification as new virus species. For example, the variant of Gadgets Gully virus identified here shares 92.7% amino acid similarity and 80.7% nucleotide similarity to the currently available genome (YP_009345034.1) originally isolated in 1976 [24]. Given the commonly applied rules for species delineation (< 90% amino acid similarity and/or < 80% nucleotide similarity), the Gadgets Gully variant detected here is the cusp of being considered a new species. Similarly, given the sequence divergence between the original Gadgets Gully virus strain (CSIRO122) isolated 1976 from *Ix. uriae* collected from Macquarie island and that collected here, and assuming typical RNA virus evolutionary rates of between 1 x 10^−3^ – 1 x 10^−4^ nucleotide substitution per site per year, it would take approximately 100–1000 years to reach that level of divergence. Although simplistic, this analysis suggests that the divergence of the Macquarie island and Neko harbour Gadgets Gully virus variants took place at least several decades ago. Conversely, one of the original Okhotskiy virus strains isolated in 1972 (ATW68806.1) and the Okhotskiy virus strain sequenced here share 97.7% amino acid similarity and 87.6% nucleotide similarity in the VP1 (RdRp) segment, indicative of a more recent common ancestry. Regardless of how and when these viruses became transferred to new locations, it is evident that there has been circumpolar dispersal of *Ix. uriae* associated viruses.

### Virus–host co-evolution and migration

To better understand virus–tick co-evolution, the host mitochondrial genome of *Ix. uriae* was mined from all sequence libraries, and a phylogenetic analysis performed with a set of representative outgroup ixodid species. For comparison, the RdRp sequence region of a subset of closely related bunyaviruses found either at the northern or the southern sampling sites was similarly subjected to phylogenetic analysis. Although our *Ix. uriae* sequences only represent a small portion of the distributional range of both circumpolar regions that this species inhabits, the results obtained are in agreement with previous studies in identifying two distinct tick populations with no evidence of dispersal between the northern and southern hemispheres (Figure 2) [26,32]. In addition, we observed longer branch lengths in the Antarctic *Ix. uriae* mitochondrial sequences than those from northern Sweden, indicating greater genetic diversity in the southern circumpolar tick distributional range. Indeed, a higher mean genetic diversity (p-distance) was observed between the mitochondrial sequences within the southern population (0.032) than the northern population (0.001). Although more data is clearly needed, this observation is compatible with the idea that the southern circumpolar population represents the ancestral population.

**Figure 2.**
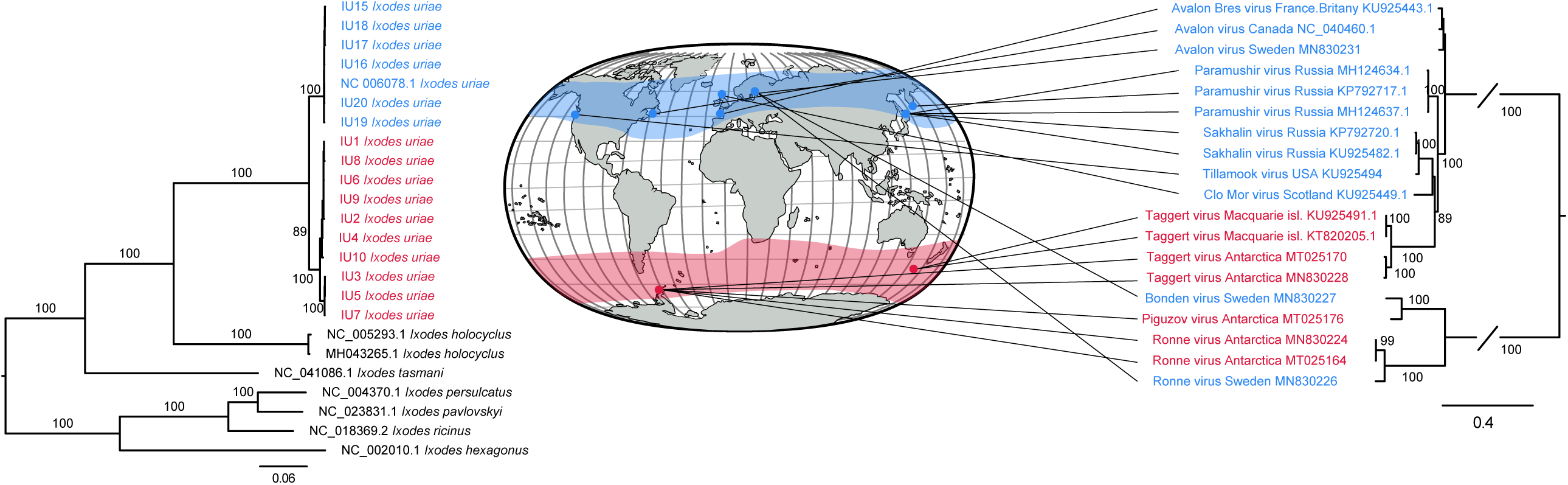
Phylogenetic analysis of near complete host mitochondrial genome data (mined from each library) and of closely related bunyaviruses found at either the southern (red colour) or northern (blue colour) circumpolar sampling sites. Lines connect virus with sampling location. Numbers on branches indicate SH support (only branches with SH support ≥80% are indicated) and branch lengths are scaled according to the number of amino acid substitutions per site. All phylogenetic trees were midpoint-rooted for clarity only.

The phylogenetic analysis of a subset of closely related bunyaviruses reveals several dispersal patterns (Figure 2). The clade that includes Avalon virus, Paramushir virus and Taggert virus indicates that *Ix. uriae*-associated viruses appear to move within the circumpolar regions inhabited by *Ix. uriae*. For example, Avalon virus, closely related to Paramushir virus [33], has been found at the Great island, Newfoundland, Canada [29], along the coast of Brittany, France [34], and now in the northern part of Sweden. As noted earlier, there are also indications of movement between the northern and southern circumpolar regions. Indeed, the close phylogenetic relationships of both Bonden virus and Ronne virus to southern circumpolar viruses clearly demonstrate that viruses are able to move between these two highly disjunct regions. In addition, that we can see such dispersal with only two sampling sites suggests that this movement is not infrequent. More generally, that there are several different bird species, many of which are known hosts for *Ix. uriae* [11], that have migratory routes within and between the circumpolar regions [35] and cover the geographical distribution of *Ix. uriae*, suggests that *Ix. uriae* associated viruses are being transported by migratory birds within and between the circumpolar regions. Hence, it is possible that these viruses may be present in all regions with permanent populations of *Ix. uriae*.

### Novel and previously identified RNA viruses

We discovered a number of other novel and previously identified RNA viruses. For example, Bulatov virus and Vovk virus (*Hepeviridae*) have previously been identified in Antarctic ticks [27] and were also discovered here (Figure 1, Table 1). They share a common ancestor and are relatively similar genetically (86.5% amino acid identity), but are in themselves divergent, sharing only 35.1% and 36.8% amino acid similarity, respectively, to the tick-borne tetravirus like virus (AII01815). Umea virus (*Mononegavirales*) was discovered in several of the tick libraries from northern Sweden and was relatively abundant (1,438 reads per million) in one library (Table 1). It shares a most recent common ancestor with Genoa virus (Figure 1) identified from *Ix. holocyclus* ticks from Australia [36], but is again relatively divergent (56.4% amino acid similarity). Similarly, Ubmeje virus (*Bunyavirales*) was identified in several libraries from northern Sweden, although it was not abundant (Table 1). Ubmeje virus shared only ∼35.8% and 35.9% amino acid identity to Bronnoya virus and Ixodes scapularis bunyavirus, both previously observed in ixodid ticks [37,38], and which together form a monophyletic group (Figure 1).

In the Antarctic sequence libraries we observed Gerbovich virus (*Picornaviridae*) that has previously been described in this region [27]. The two variants are very similar in sequence (99.7% amino acid similarity, 99.3% nucleotide similarity), but have only 56.5% amino acid similarity with their closest ancestor, Hubei tick virus 1, identified a pool of ticks from China (Figure 1) [39]. Aside from Okhotskiy virus (*Reoviridae*) described above, we also identified Fennes virus in the Antarctic sequence libraries, with near identical sequence similarity (99.8% amino acid similarity, 100% nucleotide similarity) to the sequence of this virus identified previously [27]. Fennes virus represents a highly divergent lineage, sharing only 31.0% amino acid similarity with Shelly headland virus discovered from *Ix. holocyclus* ticks from Australia [36].

Two novel orthomyxoviruses were identified: Uumaja virus from northern Sweden and Neko harbour virus from Antarctica (Figure 1, Table 1). Uumaja virus shares a common ancestor with Zambezi tick virus 1, previously identified in a *Rhipicpehalus* sp. tick collected in Mozambique [40]. Neko harbour virus was found to be abundant (more than 2,000 reads per million) in a single library (Table 1) and clusters with Aransas Bay virus [41], to which it was most similar (82.7% amino acid similarity), Jos virus [42], Thogoto thogotovirus [43] and Upolu virus [41], all of which are tick derived and originate from different continents. This pattern is indicative of a long-term association between ticks and these viruses.

Finally, we identified two novel and divergent viruses within the *Tombusviridae*: Paradise bay virus from Antarctica and Upmeje virus from the northern Sweden (Figure 1, Table 1). Paradise bay virus grouped phylogenetically (50.9% amino acid similarity) with Hypsignathus monstrosus tombus-like virus 2, a virus sequenced from blood samples of Hammer-headed fruit bats (*Hypsignathus monstrosus*) collected in the Republic of the Congo [44]. Upmeje virus was found to be highly divergent, sharing only 35.2% amino acid similarity with Sanxia water strider virus 14 and did not group with any other viruses (Figure 1). It is noteworthy that Upmeje virus was the most abundant in our study, reaching more than 10,000 reads per million, some five times higher than the host marker gene abundance in the same library (Table 1).

## Discussion

Ticks are the most important blood-feeding arthropods in temperate and polar regions [45–53]. Their unique ability to adapt to harsh, climatologically variable environments and different types of vertebrate host has enabled them to become established in many different types of habitats and parts of the world. In addition, ticks are well-known vectors of multiple viruses, including those that are known pathogens to humans and other animals, as well as viruses considered or likely to be symbionts [36,37,54]. We studied the virome of the seabird tick, *Ix. uriae*, that occurs within both circumpolar regions, as a means to understand virome composition, host–virus co-divergence and the long-distance dispersal of tick-borne viruses. In particular, we focused on whether we could infer dispersal events within and between the southern and northern circumpolar regions that are separated by a substantial geographic distance.

Following virome analysis of 16 *Ix. uriae* meta-transcriptomic libraries, we identified 16 RNA viruses of which seven were novel (Figure 1, Table 1). Of the nine viruses previously discovered, several have been documented in *Ix. uriae* and some have also been shown to have pathogenic properties to either birds and/or humans [55,56]. For example, Gadgets Gully virus was originally found on Macquarie island, south-west of New Zealand, in the 1970s [24,57], then from ticks collected there in 2002 [25], and now again here from ticks collected in 2018 at Neko harbour, Antarctica. The levels of sequence diversity between the previously collected strains and the sequence from the present study suggests that Gadgets Gully virus and likely many other viruses have been maintained in circumpolar regions for several decades, if not centuries. Similarly, we identified Avalon virus, previously found in Canada and France, in the ticks collected at Bonden island. This supports the idea that viruses are being transported within the circumpolar regions inhabited by *Ix. uriae*. These viruses could either be transported with the ticks carried by seabirds during migration or directly by infected birds. Indeed, several well-known bird-hosts of *Ix. uriae* have circumpolar migration patterns. For example, in the southern ocean, birds of the order Procellariiformes show circumpolar migration involving many stopover sites [58,59]. In the arctic region, many Charadriiform birds undertake seasonal long distance longitudinal migrations. Similarly, Black legged kittiwake (*Rissa tridactyla*), Atlantic puffin (*Fratercula arctica*) and Thick billed murre (*Uria lomvia*) all show seasonal movements between the eastern and western north Atlantic [60,61].

We also saw evidence of historic movement between the northern and southern circumpolar regions. In particular, our phylogenetic analysis revealed that Bonden virus, identified in ticks from northern Sweden, was closely related to Piguzov virus from Antarctica [27]. Similarly, Ronne virus, present in Antarctica, was also found in northern Sweden (Figure 1, Figure 2). The close evolutionary relationship of viruses from the northern and southern circumpolar regions suggests that they have moved between the poles after the *Ix. uriae* population diverged into two sub-populations. Given that the northern and southern *Ix. uriae* populations are phylogenetically distinct [26], it seems likely that it is viruses rather than the hosts that are transferred between the two polar regions. This, in turn, implies that it is virus-infected migratory birds that transport the viruses between the poles. Although some birds migrate very long distances, few species are known to move between the Arctic and Antarctic regions. The Arctic tern (*Sterna paradisea*) has the longest migration distance of any avian species [62]. Although there are no records of *Ix. uriae* found on this species, it is likely that they can become infested as they breed in dense colonies near bird species that are well-known hosts of *Ix. uriae*. As noted above, Procellariiform birds are long distance migrators and a study using geolocators of the Short-tailed shearwater (*Ardenna tenuirostris*), revealed that this species migrated to south of the Antarctic Polar Front after their breeding period in Tasmania, and following a stopover migrated northward to spend the Arctic summer in a location as far north as the Bering sea [63]. Additionally, the sooty shearwater (*Ardenna grisea*) also undertakes long distance trans-equatorial migrations [64], and other Procellariiform birds breeding in the Antarctic region, such as the Black-browed albatross (*Thalassarche melanophris*) and the South polar skua (*Stercorarius maccormicki*), have been occasionally observed in the Arctic region. As both the nymph and adult tick can feed for up to 12 days [13,65], these birds could theoretically act as vehicles for inter-polar virus spread.

Under what circumstances could a tick then be transferred between the two polar-regions? Unless a journey is made directly between the polar-regions, it would be necessary to occur sequentially both with respect to bird stop-over and tick life-stage development. For example, a nymph would initially latch onto the host shortly prior the bird migration, feed for the entire duration of the flight, and develop to the next life-stage during the stop-over. The adult tick would then need to find a new host to continue its journey. Given the long distances during which tick and virus have to survive, such events are unlikely to occur on one migration step. At the same time, that we could identify two clear cases of cross-circumpolar dispersal of closely related viruses from such a small sample of viruses indicates that cross-circumpolar transmission may not be infrequent.

The finding of several previously discovered and novel viruses that are known or likely to be tick-associated raises interesting questions about how these viruses are maintained in nature. In the northern circumpolar distribution, the tick life cycle can last up to seven years depending on host availability and temperature, and ticks may spend up to eleven months of the year off the host [13,15]. Hence, for most of their lifetime, the tick spends the majority of the time off host in tick aggregation formed in moist environments [17]. For viruses to be maintained and transmitted yearly within the tick population, it is arguable that unless the bird hosts develop a chronic viral infection that lasts for a year or these viruses are transmitted via other routes than via blood - that is, directly between hosts - the maintenance of these viruses at a particular location is to a large extent driven by tick behaviour and presence. In the case of tick-borne encephalitis virus, it is hypothesised that *Ix. ricinus* acts as both reservoir and vector for the viruses associated with this tick [66,67]. In particular, the behaviour of *Ix. uriae*, forming off-host aggregations, combined with the occasionally relatively short questing period for specific hosts [15,19,68], suggests that the tick non-viraemic co-feeding and trans-stadial transmission [48,66,69,70] of viruses are important in the establishment and maintenance of viruses in circumpolar environments. Despite the key role played by ticks, our study suggests that it is the avian host that likely functions as the dispersal agent of viruses along the circumpolar regions.

In sum, we have shown that the seabird tick *Ix. uriae* harbours an extensive diversity of viruses belonging to several different virus families and orders, and that there has been a transfer of viruses both within and between the northern and the southern circumpolar regions. As such, we stress the importance of the millions of birds that each year migrate across the globe and that have the capacity to transfer viruses to and from adjacent and distant geographical areas.

## Materials and Methods

### Tick collection and total RNA extraction

Adult female *Ix. uriae* ticks were collected during 2016–2017 from the Bonden island bird station in northern Sweden (lat/long: 63.433617, 20.038344) and from the ground around a Gentoo penguin (*Pygoscelis papua*) colony at Neko harbour, Antarctica (lat/long: - 64.824066, −62.665999), during 2018. All ticks were morphologically keyed to species [71] and were subsequently stored in −80°C until further processing. Prior to total RNA extraction, ticks were washed in PBS buffer two times and then pooled (Supplementary Table S1). Total RNA from 16 tick pools was then extracted using the RNeasy® Plus Universal kit (Qiagen) following the manufacturer’s instructions.

### Sequence library construction and sequencing

Sequencing libraries, data generation and analysis was performed as previously described [37,72]. Briefly, ribosomal RNA (rRNA) was depleted from the total RNA extracts using the Ribo-Zero Gold (human-mouse-rat) kit (Illumina) following the manufacturer’s instructions. RNA sequencing libraries were then prepared for all rRNA depleted extracts using the TruSeq total RNA library preparation protocol (Illumina) followed by paired-end (150 bp read-length) sequencing on a single Illumina HiSeq X10 lane, as performed by the Beijing Genomics Institute, Hong Kong. The raw sequence output was then quality trimmed with Trimmomatic v.0.36 [73] using the default settings for paired-end sequence data and assembled *de novo* using Trinity v.2.5.4 [74] with read normalisation apart from default options.

### Virome analysis and presence across libraries

All *de novo* assembled contigs were initially screened against the complete non-redundant nucleotide and protein databases (NCBI GenBank) using blastn v.2.6.0+ [75] and Diamond v.0.9.15.116 [76], respectively, employing cut-off e-values of 1 × 10^−5^ for both methods. To further assess the data and to identify potential endogenous viral elements, all assemblies indicative of being of RNA virus origin were screened using the NCBI Conserved Doman Database (www.ncbi.nlm.nih.gov/Structure/cdd/cdd.shtml) with an expected value threshold of 1 × 10^−3^. Relative abundance of the identified viruses was determined by comparing the mapping results of the *Ix. uriae* mitochondrial cytochrome C oxidase I (COX1) gene (NC_006078.1, positions 1214–2758) against all RdRp containing contigs using Bowtie2 v.2.3.4 [77], employing the default local setting for all libraries. Relative abundance was calculated as reads per million: that is, the number of reads mapped to a contig divided by the total amount of reads in a library multiplied by a million. A particular virus was considered abundant if (i) it represented >0.1% of total ribosomal-depleted RNA reads in the library, equivalent to a reads per million value of 1000 or more, and (ii) if the abundance was higher than that of the host COX1 gene [37,72]. If the relative abundance of a virus contig was less than 1 read per million mapped, or below the level of cross-library contamination due to index-hopping measured at 0.1% of the most abundant library for the respective virus species, the library was considered negative for the presence of the virus contig. A virus was considered novel if the RdRp region showed < 90% amino acid or < 80% nucleotide similarity to any previously identified virus.

### Virus evolutionary history

To infer the evolutionary history of all the RNA viruses identified here they were combined with representative amino acid data sets of the RNA-dependent RNA-polymerase genes of viruses from the orders *Bunyavirales, Mononegavirales, Orthomyxovirales*, the families *Picornaviridae, Reoviridae*, the Hepe-Nido-like and Tombus-Noda-like groups and the genus *Flavivirus*. These sequences were then aligned using the E-INS-i algorithm in Mafft v.7.271 [78]. To reduce alignment uncertainty, regions that aligned poorly were removed using TrimAl v. v1.4.rev15 [79] under the ‘strict’ settings. Each alignment was then subjected to model testing to determine the best-fit model of amino acid substitution using ModelFinder [80] via IQ-TREE v.1.6.12 [81]. Finally, maximum likelihood phylogenetic trees of each data set were estimated using the IQ-TREE package, implementing a stochastic hill-climbing nearest-neighbour interchange tree search. Phylogenetic robustness was assessed using Shimodaira–Hasegawa (SH)-like branch supports.

### Virus–tick evolutionary history

To compare the evolutionary history of *Ix. uriae* and a subset of closely related *Bunyavirales* found at both sampling sites (see Results), the mitochondrial genome of *Ix. uriae* (NC_006078.1) was used as reference for mapping with Bowtie2 v.2.3.4 [77], with local settings, against all sequence libraries. The resultant mitochondrial nucleotide consensus sequences from each sequence library were then aligned with a set of Ixodidae reference sequences using the G-INS-i algorithm in Mafft v.7.271 [78]. Following visual alignment checking in AliView v.1.26 [82], model testing and estimation of a maximum likelihood phylogenetic tree was performed in IQ-TREE as described above. The corresponding virus phylogeny was inferred using the nucleotide RdRp open reading frame sequences of a subset of bunyaviruses, keeping the open reading frame intact and utilising the same model testing and phylogenetic tree inference procedure as described above. All phylogenetic trees computed were visually compared and edited with FigTree v.1.4.3 (https://github.com/rambaut/figtree/). Comparison of genetic diversity between the northern and southern populations was undertaken by computing the number of base differences per site averaged over all sequence pairs between the two populations (i.e. p-distances), via Mega X v.10.1.1 [83], using including transitions and transversions, uniform rates, pairwise deletion of gaps and 500 bootstrap replicates to estimate variance.

## Data availability

The raw sequence data generated here have been deposited in the NCBI Sequence Read Archive (BioProject: PRJNA594749) and all virus related sequences have been deposited on NCBI GenBank (accession numbers: MN830224–MN830247).

## Acknowledgments

The authors thank Thomas G.T. Jaenson (Uppsala University) for insightful and constructive comments that improved the manuscript significantly. J.H.O.P is funded by the Swedish research council FORMAS (grant nr: 2015-710). E.C.H is funded by the Australian Research Council, Australian Laureate Fellowship (FL170100022).

## Supplementary Information

**Supplementary table S1**. Sequence library information regarding sampling location, host species and collection year.

**Supplementary table S2**. Read mapping results for each identified virus and host gene per individual library.

